# Establishing parallel and scalar measures of executive and semantic demand: an online norming study of matched, cross-domain judgement tasks

**DOI:** 10.64898/2026.09.24.753760

**Authors:** Georgia Bozianu, Alex Woolgar, Matthew A. Lambon Ralph, Ajay Halai

## Abstract

Task demand has a long-standing history in experimental psychology, with numerous investigations of the capacity of different systems and processes under varying cognitive load. Whilst demand is a broadly used term in the literature, it is limited by a lack of cross-domain operationalisation, and thus can be a confound in many inter-domain investigations. To address this, we generated and validated two - one semantic and one visuospatial - large-scale, open-source, widely applicable stimuli sets that: (i) can probe a wide, continuous range of objective behaviourally-measured processing demand, and (ii) can be directly matched across two highly contrastive cognitive domains - here semantic and visuospatial tasks used as test beds. We normed these tasks in a large, online cohort of healthy adults, enabling future studies to utilise and benefit from reliable baseline measures of demand. Using a matching algorithm that takes into account both speed and accuracy, we then closely paired each semantic judgement trial with a trial of comparable difficulty from the corresponding visuospatial task.to produce an open-source, cross-domain, matched set of over 400 items. Finally, we analysed the psycholinguistic and perceptual properties of these stimuli to explore some of the underlying factors driving demand.

## Introduction

Cognitive operations are often described as a series of computations that transform input into output via an intermediate neural process (Bechtel, 2008b, 2008a; Roediger III et al., 1999). Across domains, these operations are influenced by the level of demand imposed on the system: performance typically slows and ultimately deteriorates as stimuli, tasks or responses become more difficult or ambiguous. Such effects are observable in language and other higher-order cognitive processes, and are sometimes explored or manipulated in cognition paradigms across different study populations and clinical groups (Becker et al., 2016; Earles et al., 2004; Huber, 1985; Robinson, 2001; Snowden et al., 2001). Unless the goal is threshold detection (as often is the case for psychophysics experiments), it is rare for experiments to consider the full range of performance, spanning from straightforward to very difficult trials. Indeed, in contrast to the tradition of systems evaluation in engineering (where the full range of operation demand is checked against the design brief), sensory and cognitive processing tends to focus on finding the limits of the target domain. Moreover, despite the ubiquity of research into it, cognitive demand often remains a broad and underspecified construct, often inferred rather than directly measured.

Variation in demand can be considered at multiple stages of a computation, including in relation to the quality or complexity of the input (Iyer et al., 2022; Küper & Karbach, 2016; Seyderhelm & Blackmore, 2023; Wasow et al., 2003), the efficiency of internal processing (Achard & Bullmore, 2007; Curtin et al., 2019; Duering et al., 2013; Marinho et al., 2019), or the constraints imposed on the output (Jahfari et al., 2010; Orthey et al., 2019; Ross et al., 2025). However, not all forms of demand are explored in the same way. Sometimes difficulty is intentionally engineered through experimental design or task structure, whereas in other contexts it may arise naturally from properties inherent to the stimuli themselves. Thus, in the present work, we differentiate between intrinsic and extrinsic demand.

Intrinsic demand refers to difficulty that is linked to the stimulus itself - properties that are more stable across contexts and are less dependent on task instructions or response requirement. In language processing, psycholinguistic properties, including word frequency or length, reliably influence processing effort across tasks and populations (Cohen et al., 2021; Monsell et al., 1989; Pallotti, 2019; Toyama, 2021). Analogue intrinsic properties exist outside the linguistic domain, such as the complexity of a visual array, the dimensionality of a spatial configuration, or the computational load of a mental arithmetic problem.

In contrast, extrinsic demand arises from factors external to the stimulus, such as the number of response options, the number of rules governing a task, or the requirement to switch between tasks sets (Bindemann et al., 2005; Reisenauer & Dreisbach, 2014; Verbruggen et al., 2004). These manipulations are typically imposed top-down by the experimenter and can be adjusted to fine tune task difficulty. While extrinsic demand plays a critical role in cognitive control paradigms, it is conceptually distinct from intrinsic demand in that it does not reflect stable properties of the stimuli themselves.

Although intrinsic demand is widely assumed to influence behaviour, it is rarely quantified in a systematic manner. In language research, stimulus difficulty is often approximated using individual proxies (e.g., word frequency, length, imageability, etc), while non-linguistic tasks rely on bespoke manipulations tailored to specific paradigms. As a result, comparing across domains is challenging unless task difficulty is equated (see Humphreys & Lambon Ralph, 2017) and the comparisons are limited to two levels of difficulty (e.g., easy and hard trials). Critically, there is currently no unified, empirically grounded scale that allows for intrinsic demand to be compared across domains using matched tasks and common behavioural metrics across a range of performance that spans from different levels of accurate performance to the “breaking point” (i.e., where slowed response times are joined with increased error rates).

This limitation has important consequences for both basic and applied research. In neuroimaging and neurostimulation studies, changes in neural recruitment are frequently interpreted as reflecting increased cognitive demand (Duncan, 2001, 2010; Hallam et al., 2016; Martin et al., 2023; Webler et al., 2022; Woolgar et al., 2011, 2015), yet without a wealth of cross-domain, normed information it is difficult to determine whether such changes reflect individual differences within a population or other sources of variation such as stimulus-driven difficulty, task-specific strategies, or stimulation effects. Similarly, in clinical research, particularly in the context of stroke or other acquired brain injuries, adaptive responses to increased demand have been extensively studied across language, motor, and attentional domains (El Hachioui et al., 2013; Gallucci et al., 2024; Lomas & Kertesz, 1978; Saur et al., 2006; Wilson et al., 2023; Woolgar et al., 2013; Yan et al., 2025; Zhao et al., 2018). However, the interpretation of such findings is constrained by the absence of a comprehensive characterisation of how demand impacts cognitive domains across a wide range of performance levels, including beyond a breaking point.

Addressing these issues requires new systematically-collected normative data that evaluates the full scalar range of intrinsic demand across contrastive cognitive domains. Such a resource would allow stimulus difficulty to be quantified independently of task structure, provide a common scale for comparing different domains, and offer a principled basis for designing behavioural and functional neuroimaging investigations in neurotypical and clinical populations. The present study addressed this gap by developing a formal measurement scale to operationalise intrinsic demand across two domains: semantics and visuospatial processing. Their distinct cognitive architectures allow us to differentiate between domain specific architectures and shared systems, which would thus provide strong evidence for the role of domain general mechanisms. Moreover, their prevalence in the literatures surrounding control mechanisms (both specialised and general) provide a rich foundation of knowledge from which to build upon our understanding of how increasing demand affects these domains. We collected behavioural data from a healthy adult population to capture graded variation in task difficulty for semantic and visuospatial processing, by systematically sampling across the full range of easy to challenging trials and measuring performance using response time and accuracy.

## Methods

### Participants

We recruited participants via Prolific (www.prolific.com) to complete the study online and reimbursed them at a rate of at least £6 per hour. Eligibility criteria required participants to be UK residents and nationals, native English speakers, aged 18–40 years, with no self-reported literacy difficulties or language-related disorders, and normal or corrected-to-normal vision.

A total of 152 participants were recruited across both tasks. Fifty participants completed the Semantic Judgement Task (SJT; 454 trials). The Visuospatial Grid Task (VGT) comprised two independent stimulus sets of 500 trials each, which were completed by two separate groups of 51 participants (102 participants total). All participants provided informed consent prior to participation.

### Stimuli

We adapted the SJT from a previously validated synonym judgement test (Jefferies et al., 2009). On each trial, participants were presented with a probe word and were required to select, from three alternatives, the word most closely-related in meaning to the probe. To create an expanded stimulus set, we randomly selected probe words (nouns and verbs) from the English Lexicon Project (Balota et al., 2007). Candidate probes were sampled to span a broad range of psycholinguistic properties, including word frequency, imageability, and syllabic length, with distributions designed to approximate those observed in English more generally. For each probe, a synonym/semantically-close target and two unrelated distractors were selected and matched to the probe on syllabic length, word frequency, and imageability. To verify that the semantic distance between the probe and the target on a given trial was smaller (i.e. the words had more similar meanings) than the semantic distance between the probe and the distractors, we compared measures of semantic distance (derived from the glove-wiki-gigaword-300 embedding set: Pennington et al., 2014) between Probe-Target (PT) and Probe-Distractor (PD) word pairs using Bayesian paired-samples *t*-tests. The analysis provided extreme evidence that PT scores were smaller than the mean PD per trial (*M* difference = - 0.225, *t*(448) = -26.09, BF₁₀ = 1.55 × 10⁸⁸, *d* = 1.54), as well as both individual PD semantic distance scores (Distractor 1: *M* difference = -0.227, *t*(448) = -23.84, BF₁₀ = 1.16 × 10⁷⁸, *d* = 1.48; Distractor 2: *M* difference = -0.223, *t*(448) = -25.28, BF₁₀ = 3.64 × 10⁸⁴, *d* = 1.45). We provide full details of the stimulus selection procedure in the Supplementary Materials: 1.

The VGT comprised 1,000 checkerboard-style grid stimuli generated in MATLAB (MathWorks Inc., 2022). To mirror the form of the SJT, each trial consisted of a probe grid and three comparison grids, one of which was identical to the probe, excepting some potential spatial transformations in harder trials. Grid stimuli varied systematically along five dimensions: grid size (3×3 or 4×4), number of colours (two or three), colour distribution (approximately equal or unequal), rotation (0° or 90/180/270°), and flipping (horizontal or vertical plane). Target grids were exact copies of the probe grid, sometimes rotated and/or flipped, while distractor grids additionally differed from the probe by the colour of a single cell. We designed these manipulations to sample a range of visuospatial complexity while maintaining a systematic task structure across trials. See Figure 1 for examples of both easy and hard grid and semantic trials. We provide further details of grid construction in the Supplementary Materials: 2.

**Figure 1:**
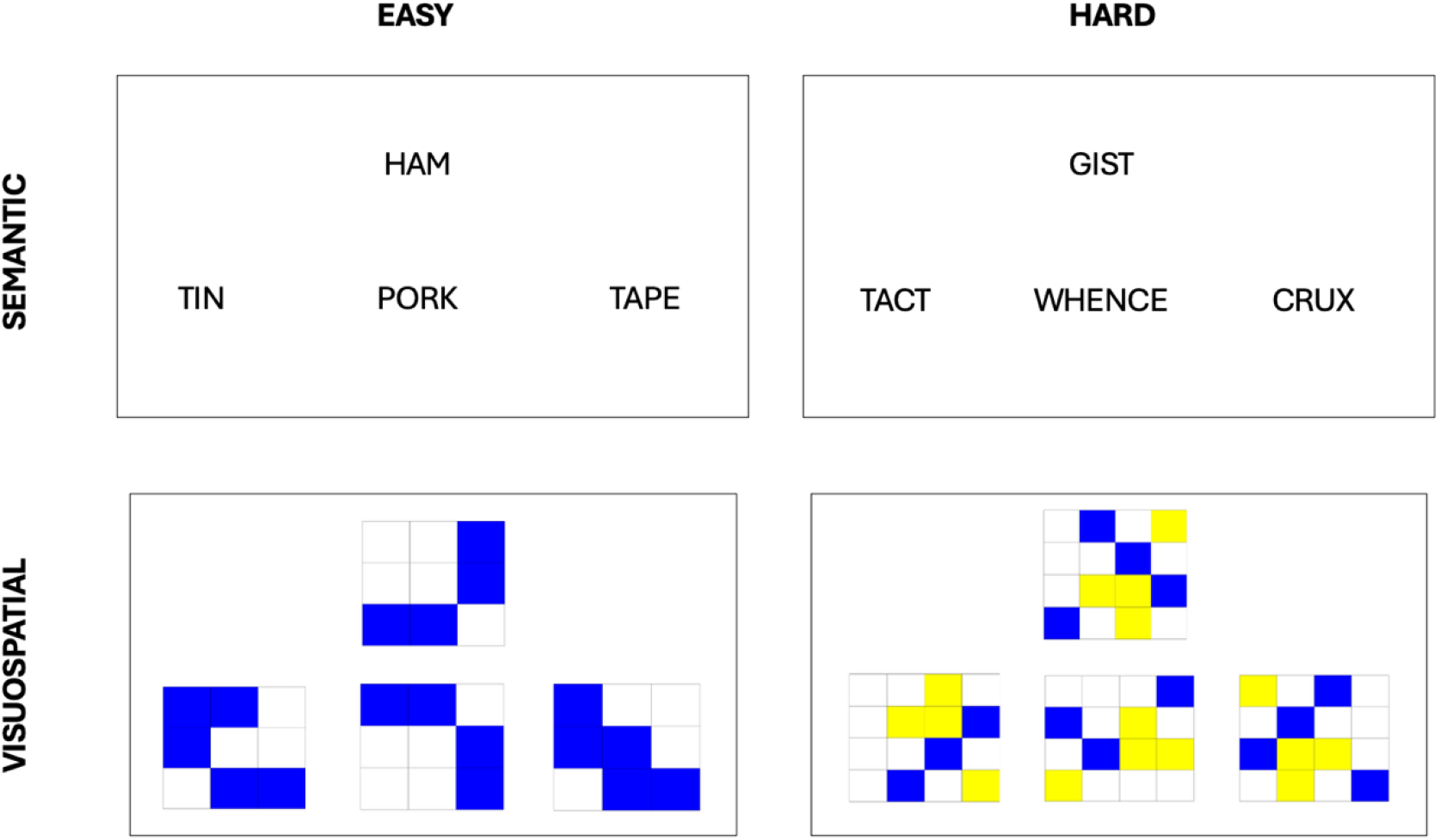
Trial Examples from Semantic Judgement Task (SJT) and Visuospatial Grid Task (VGT)

### Procedure

Both tasks employed a three-alternative forced-choice paradigm. Each trial began with a fixation cross presented for 500ms. The probe stimulus was then displayed at the top of the screen with three stimuli options presented simultaneously below. Participants were instructed to select the option that was the most semantically (SJT) or visually (VGT) similar to the probe by pressing the key 1, 2, or 3 on the keyboard, which corresponded to the left, centre, or right stimulus option. Trials were presented in a fully randomised order, and the location of targets and distractors was randomised across participants. Responses were permitted within a 4500ms window, after which time the next fixation cross began. If no response key was received within the 4500ms time window, that trial was recorded as incorrect. The experiment was self-paced, and participants were offered two optional one-minute breaks. Prior to the main experiment, participants completed at least one round of practice trials with feedback (N=15 across maximum of two rounds) and were required to achieve at least 75% accuracy to proceed.

All tasks were implemented using jsPsych and hosted on the MRC Cognition and Brain Sciences Unit’s JATOS server (Lange et al., 2015). Task scripts and stimulus generation code can be found at the Open Science Framework (https://osf.io/kfdj3/overview) or on GitHub (https://github.com/georgiabozianu/crossdomain-matched-norming/tree/main).

### Data Processing & Analyses

Primary behavioural measures were response time and accuracy. We scored trials as incorrect if participants selected one of the two distractors instead of the target or if they failed to respond within the allotted 4500ms time window. We screened data for evidence of non-compliant responding, defined a priori as a) repeated execution of the same response on five or more consecutive trials (including non-responding); b) the repeated occurrence of a systematic response sequence on three or more consecutive occasions; c) any occurrence of a mistaken button press (e.g. using any key other than 1, 2, or 3). These patterns were interpreted as indicating sustained inattentive or invalid responding rather than genuine task performance. We excluded trials from participants exhibiting such patterns prior to analysis and then calculated the mean response time per item using correct responses only.

We then paired trials across the two tasks using the MATCH program (van Casteren & Davis, 2007) to create a set of stimuli that are behaviourally equated across cognitive domains. The programme matched all semantic trials to a corresponding visuospatial trial using their item-wise mean response time and accuracy scores. The algorithm recursively searches the space of possible pairings to identify the solution that best equates the distributions (mean, median, and standard deviation) of both accuracy and response time across the two stimulus sets, producing closely comparable items across all levels of demand.

To identify the stimulus properties contributing to semantic and visuospatial task demand, we analysed trial-wise performance using linear mixed-effects models (LMEM) for response time and binomial generalised linear mixed-effects models (GLMEM) for accuracy. First, we selected a set of psycholinguistic variables (taken from the English Lexicon Project: https://openlexicon.fr/datasets-info/EnglishLexiconProject/README-ELP.html) for each of the probe words in the SJT. The following nine variables of interest were available for at least 429/454 probe words from the SJT: word length, syllabic length, frequency, orthographic and phonological neighbours, concreteness, semantic neighbourhood density, age of acquisition, and semantic diversity. We used a principal component analysis (PCA) to reduce these nine correlated variables into a smaller set of orthogonal components that captured the most variance across the semantic stimuli. Bartlett’s Test of Sphericity revealed that the data was suitable for factor analysis (X^2^ (66) = 130,051.68, *p* < .001). To investigate the potential driving factors of semantic demand, we then used the principal components derived from this PCA as the fixed effects in the semantic LMEM, with participant age and sex as covariates. For the VGT models, fixed effects comprised the five manipulated visual grid properties together with participant age and sex as covariates. We included subject as a random intercept to account for repeated observations within participants. Models were fit using statsmodel in python, and statistical significance of fixed effects was assessed using F-tests with Satterthwaite approximated degrees of freedom.

## Results

### Participant Information

Demographic information for participants in the SJT (n = 50) and VGT groups (n = 102) were similar. The gender (F:M) split across the two groups was 32:17 and 61:34 (1 participant in the SJT and 5 participants in the VGT preferred not to say), and the mean age (SD) was 29.24 (5.62) and 30.36 (5.88), respectively. A Bayesian contingency-table test was conducted to assess differences in sex distribution between groups. The results provided strong evidence for the null hypothesis (BF₁₀ = 0.056), suggesting that sex composition did not differ meaningfully between groups and was unlikely to represent a confounding factor. To similarly compare age differences across the two groups, a Bayesian independent-samples *t*-test indicated that the data were approximately three times more likely under the null hypothesis than the alternative (*t* = -1.12, BF₁₀ = 0.33, *d* = 0.19), providing evidence against a substantial age difference between groups.

### Data Quality Screening

All data were screened for inattentive or invalid response patterns prior to analysis. In the SJT, 329 trials (1.4% of total trials) across 18 participants were excluded due to suspicious response behaviour (e.g., unrealistically fast responses or sustained non-response). However, all 50 participants contributed valid data to the final dataset, yielding 22,371 analysable trials.

In the VGT, 408 individual trials were excluded across 43 participants due to suspicious response behaviour. Additionally, four participants exhibited extensive invalid responding (>300 trials each) and were excluded entirely from analysis. Therefore, a total of 4.8% of all trials were excluded and the final VGT dataset comprised 96 participants contributing 47,592 valid trials. Full exclusion details are provided in the Supplementary Materials: 3.

### Behavioural Performance

No participant performed below chance (33.3%) on either task. To assess behavioural performance across the two cognitive domains, for each task type (SJT and VGT), we calculated the mean response time (RT) and accuracy per trial, across participants. Trials across both tasks captured the range of performance, from fast and highly accurate through to slower and more effortful (see Figure 2, panel A). We then created a matched set (see Figure 2, panel B) by selecting the 410 SJT trials with the highest response-agreement across participants and paired each trial with a VGT trial, using the MATCH program (van Casteren & Davis, 2007). This algorithm optimally paired trials with equivalent difficulty, using behavioural measures of performance (both accuracy and response time). The matching worked as expected. Bayesian independent-samples *t*-tests were conducted to assess whether the matched word and grid sets differed in behavioural performance. Response times did not differ between conditions, with the data providing moderate evidence in favour of the null hypothesis (*M* difference = -44.09 ms, *t* = -1.70, BF₁₀ = 0.323, *d* = 0.12). Accuracy showed strong evidence for no difference between conditions (*M* difference = -0.33%, *t* = -0.39, BF₁₀ = 0.084, *d* = 0.03). Bayesian comparisons of variability further indicated no meaningful differences in dispersion between word and grid trials: posterior variance ratios were close to 1 for both response times (median ratio = 1.05, 95% CI [0.86, 1.27]) and accuracy (median ratio = 1.14, 95% CI [0.94, 1.38]). Together, these results provide evidence that the trial sets were well matched in both central tendency and variability.

**Figure 2:**
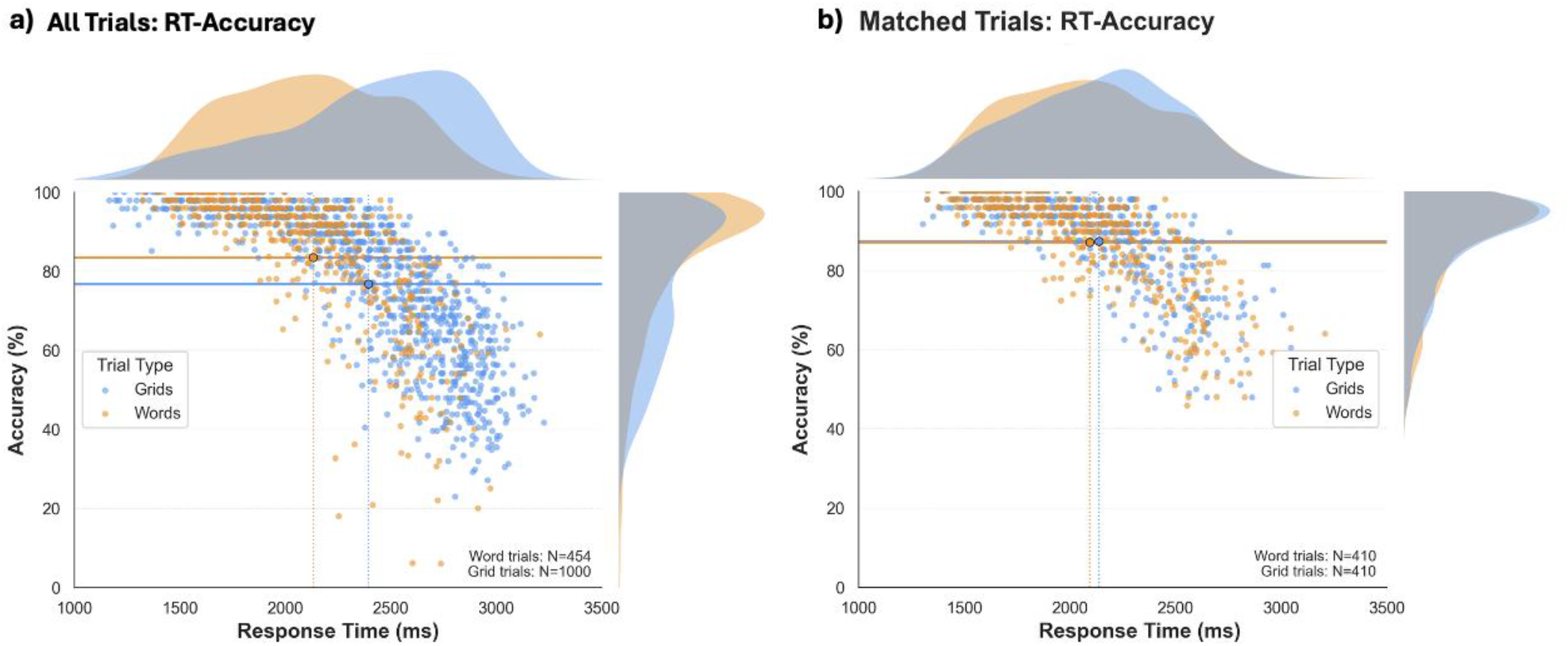
Scatter and density plots depicting the distributions of speed and accuracy per trial in the SJT and VGT pre- and post-matching. **(a)**: depicts the accuracy and response time across all trials from both tasks, pre-matching. Each data point represents a single trial, from either task. **(b)**: depicts the same measures for the filtered subset of trial pairs, after they had been optimally matched across domains.

A linear regression model examining the relationship between response time and accuracy, with task as a moderator (*mean RT ∼ mean accuracy * task),* revealed no significant interaction between task and accuracy (β = -90, p = 0.34), suggesting that the slope relating accuracy to RT did not differ across tasks. As expected, accuracy was negatively related to RT (β = -1576, p < 0.001), indicating that faster responses were generally more accurate.

### Exploratory Analyses

Participant sex significantly predicted response time (**F**(1,44.97) = 8.29, *P* = 0.006), with male participants responding more slowly than female participants (β= 225.72 ms, SE = 78.38, *t*(44.97) = 2.88, *P* = 0.006). Age was not associated with response time (β= 1.94 ms/year, SE = 6.67, *t*(44.98) = 0.29, *P* = 0.773). Neither age nor sex significantly predicted accuracy.

A varimax rotated principal component analysis on the psycholinguistic metrics revealed a three-factor solution accounting for 80.2% of the total variance in the probe word dataset. The retained components (and their relative proportion of variance explained) broadly reflected: 1) Word-Form Simplicity (47.6%); 2) Semantic Abstractness (20.9%); and 3) Lexical Familiarity (11.7%). Figure 3 shows the loadings for each task across all components. These three components were used to index intrinsic linguistic properties and were then included in mixed effects models predicting response time and accuracy in the SJT. The raw correlation matrix between psycholinguistic variables is provided in Supplementary Materials: 3.

**Figure 3.**
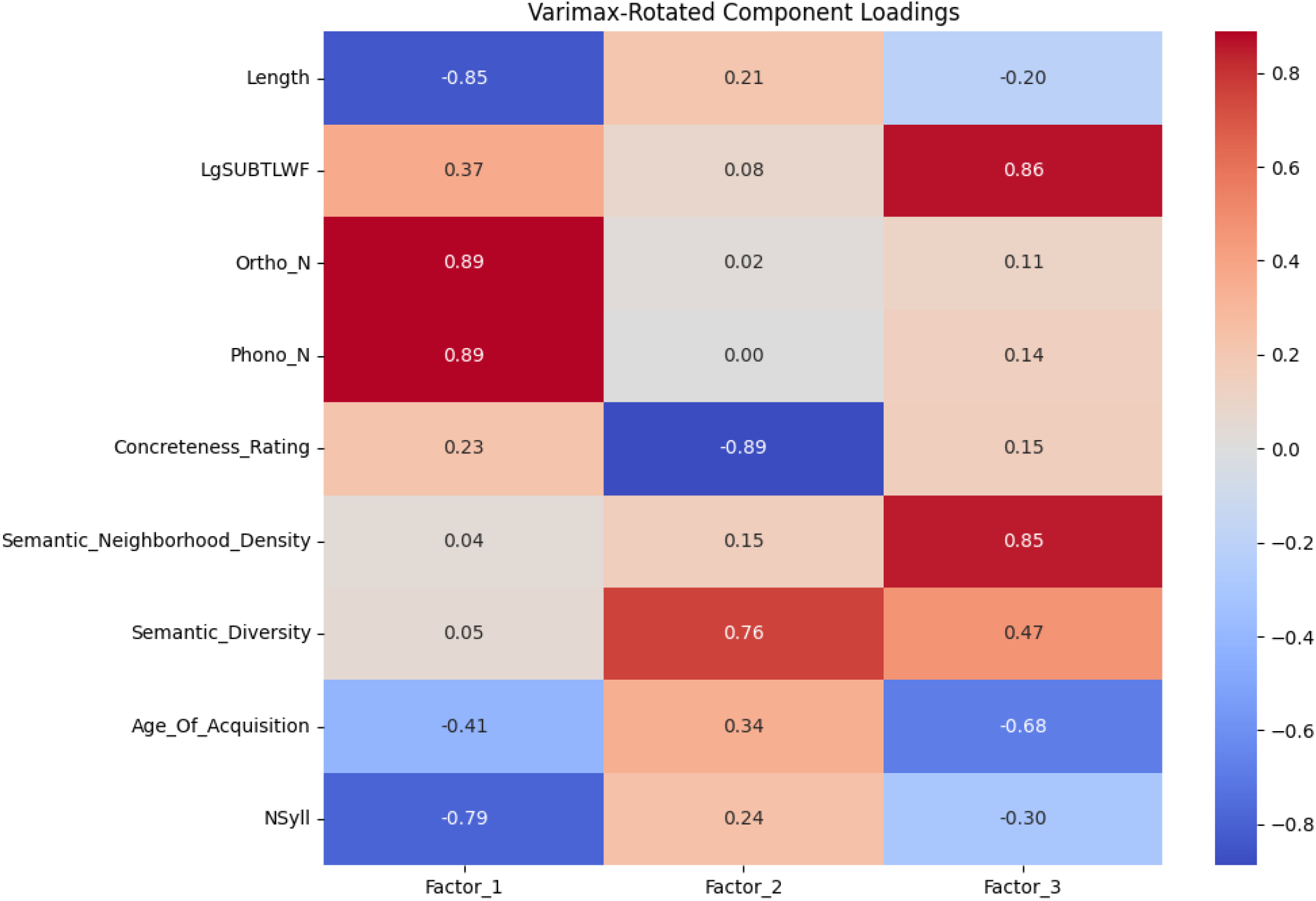
Loading plot for a varimax rotated principal component analysis on psycholinguistic variables. LgSUBTLWF = log10 transform of the SUBTLEX frequency per million words; Ortho_N = number of orthographic neighbours; Phono_N = number of phonological neighbours (excluding homophones); NSyll = number of syllables. Definitions and details of all psycholinguistic measures included in the PCA are available at the English Lexicon Project: https://elexicon.wustl.edu.

The SJT linear mixed-effects models revealed significant effects of all three principal components on response time. Factor 1 (Word-Form Simplicity) was associated with faster responses (*β* = −65.80, *SE* = 5.11, *z* = −12.89, *p* < .001), whereas Factor 2 (Semantic Abstractness) was associated with slower responses (*β* = 171.23, *SE* = 5.04, *z* = 34.00, *p* < .001). Factor 3 (Lexical Familiarity) was also associated with faster responses (*β* = −109.28, *SE* = 5.28, *z* = −20.68, *p* < .001), suggesting that participants are slower to respond to trials when the probe word is longer, has increased lexical competition, is more abstract, and less frequent. The parallel binomial generalized linear mixed-effects model predicting trial-level accuracy exhibited a similar pattern, in that Factor 2 (Semantic Abstractness) was associated with reduced accuracy (*β* = −0.471, *SD* = 0.020), and Factor 3 (Lexical Familiarity) was associated with increased accuracy (*β* = 0.319, *SD* = 0.018). The effect of Factor 1 on accuracy was small (*β* = 0.031, *SD* = 0.019), indicating little evidence for an association. A visual summary of these effects for both response time and accuracy are shown in the Supplementary Materials: 4.

In the VGT, intrinsic perceptual characteristics of the checkerboard stimuli reliably modulated task performance. Age and gender did not significantly predict VGT accuracy. Larger grid size (odds ratio [OR] = 0.85), increased number of colours (OR = 0.85), and spatial transformations (rotation [OR = 0.96] or flipping [OR = 0.97)) were associated with reduced accuracy. An unequal ratio of colours was only slightly related to higher accuracy (OR = 1.04). Further, all manipulations of grid complexity significantly increased response time: grid size (β = 508.41, *t* = 68.16); number of colours (β = 528.12, *t* = 70.24); ratio of colours (β = −121.01, *t* = −15.77); rotation (*β* = −174.17, *t* = −23.43); and flipping (*β* = −95.86, *t* = −13.04). A visual summary of the effects of these grid characteristics on both accuracy and response time is shown in the Supplementary Materials: 5.

## Discussion

The concept of cognitive demand is complex, given that many intrinsic and extrinsic factors can affect performance and the underlying cognitive and neural systems. Cognitive demand affects almost all forms of cognition, such as language, visuospatial processing, memory, attention, and executive control, among others (Archibald, 2013; de Voogd et al., 2018; Liang et al., 2017; Maujean et al., 2003; McGuire & Botvinick, 2010; Mohammadian et al., 2022). Traditionally, demand has been investigated in each domain separately; however, there is a growing interest in understanding demand – and the contributions of domain general networks – across tasks more broadly (Assem et al., 2020; Chiou et al., 2022; Darby et al., 2017; Duncan, 2010; Humphreys & Lambon Ralph, 2017; Obermeyer et al., 2020; Viviani et al., 2025). To achieve a full and equated investigation across domains, we first need to develop and validate a stimulus set that samples across a broad and continuous range of cognitive demand across the contrastive domains of interest (here using semantic and visuospatial judgements as a central test case). This normative process ensures that the relative scale of demand is explicitly defined. The behavioural measures of response time and accuracy provided objective indices of trial-level demand across semantic and visuospatial processing, enabling systematic and objective matching of items both within and across tasks.

The analyses confirmed that the manipulations designed to increase computational difficulty systematically reduced accuracy and prolonged response times. In the visuospatial grid task, increases in grid size, colour variation, and spatial transformation were associated with predictable increases in behavioural cost. In the semantic judgement task, psycholinguistic properties including word length, lexical frequency, concreteness, and semantic richness similarly modulated performance on the judgement task. These findings indicate that both stimulus sets successfully capture graded variations in task demand and thus captured the full continuous graded scale of difficulty.

In our study, demand was treated as a downstream behavioural construct of a given stimulus’ visual or semantic properties rather than being defined *a priori* in theory-specific terms. By indexing trial difficulty through observed performance, the present dataset provides a flexible framework for researchers who may differ in how they conceptualise the mechanisms underlying semantic and visuospatial demand. This approach also allows cross-domain matching without requiring strong assumptions about the precise cognitive processes engaged by individual stimuli and trials. A further strength of the dataset is its scale and internal structure. Items were sampled to achieve broad distributional coverage across relevant stimulus dimensions, allowing researchers to select subsets matched for overall demand while varying intrinsic task characteristics. The inclusion of item-level psycholinguistic and structural properties also enables secondary analyses examining specific contributors to inter-and intra-individual performance variability.

While semantic and visuospatial tasks share a common response format, inherent domain differences remain. Semantic trials may be constrained by prior knowledge and vocabulary size, whereas all visuospatial trials are theoretically solvable given sufficient time. Such differences may influence response dynamics, particularly at higher levels of difficulty. Nonetheless, the consistent behavioural scaling observed across both tasks supports their utility as comparable indices of graded cognitive demand.

One limitation of the present norms is that the study was conducted online, and, although data screening procedures were implemented to minimise inattentive responding, laboratory-based replication would further strengthen confidence in the measures. Due to the volume of data that was collected across participants, we used a between-subjects design; however, implementing a within-subjects design would allow us to minimise subject variability across tasks. Additionally, behavioural metrics provide indirect indices of cognitive effort and do not directly index underlying neural resource allocation. This means we cannot make strong claims about the processes which might dictate the speed of a participant’s response and, thus, that there are other valid ways to operationalise demand. For example, taking a long time to read a short but abstract word may be internally more effortful or require more neural resources than taking the same amount of time to read a longer but concrete and highly frequent word. However, future research may combine these stimuli with neuroimaging, electrophysiological, or pupillometric measures to examine how graded behavioural demand corresponds to neural and physiological indices of effort. The availability of large, parametrically varied stimulus sets facilitates such extensions. We propose that using this well-characterised stimulus set would allow for future work to understand more explicitly how demand is estimated in both healthy and patient populations, such as stroke and dementia. By providing carefully normed stimuli spanning a continuous demand range, the present resource enables more precise, granular, and controlled investigation of both domain-specific and domain-general cognitive processes.

## Declarations

### Ethics Approval

This study was conducted in accordance with the ethical standards of the University of Cambridge Psychology Research Ethics Committee (REC ref: 04/Q1405/66; IRAS ID: 253054).

### Consent to Participate and for Publication

All participants provided fully informed consent to take part in this study and were given the option to quit or exit the experiment at any point throughout. They were also made aware of our intention to publish the results of the study, including their data, and were given information as to the data protection and anonymity policies at the CBU before consenting to participate.

### Availability of Data and Materials

Stimuli, raw datasets, and other relevant materials are available on OSF at: https://osf.io/kfdj3/overview. Neither of the reported studies were preregistered. The scripts for data cleaning and analysis used during the current study are available on GITHUB in the cross domain-matched-norming repository, [https://github.com/georgiabozianu/crossdomain-matched-norming/tree/main].

### Open Practices

All materials and code relevant to the present study are available at OSF and GITHUB (linked above). Neither of the reported studies were preregistered.

