## Supplementary Document for "Establishing parallel and scalar measures of executive and semantic demand: an online norming study of matched, cross-domain judgement tasks"

### Supplementary Materials

Supplementary Materials 1.

To extend the 96-item test set validated in previous patient work (Jefferies et al., 2009), an additional 500-item probe list was randomly generated. Each probe was entered into Google's search engine synonym generator ('Google English Dictionary' whose data is derived from Oxford Languages: https://languages.oup.com/google-dictionary-en/) to generate related words. The top five words were then selected as candidates; the target word was selected from these candidates based on how well-matched its syllabic length, word frequency, and imageability score was to the probe. If a word did not have an appropriately related target candidate, it was discarded from the stimuli set. Once all probe words had a target word, two random, unrelated distractors were generated using the LexOps software (https://jackedtaylor.github.io/LexOPSdocs/lexops-shiny-app.html), to ensure close matching on the same psycholinguistic variables. This process was repeated until all probe words were matched with suitable targets and distractors or discarded if this failed.

Supplementary Materials 2.

Grids were constructed in MATLAB (R2023b; MathWorks Inc) to systematically sample across the hypothesised demand space for checkerboard-style stimuli. All trials had the basic manipulations (dimension, colour, ratio) and half of the trials also had the rotation and/or flip manipulation applied to *only* the target and distractor stimuli (not the probe). Source code available at https://github.com/georgiabozianu/crossdomain-matched-norming/tree/main.

Supplementary Materials 3*: Correlation Matrix of Psycholinguistic Variables*

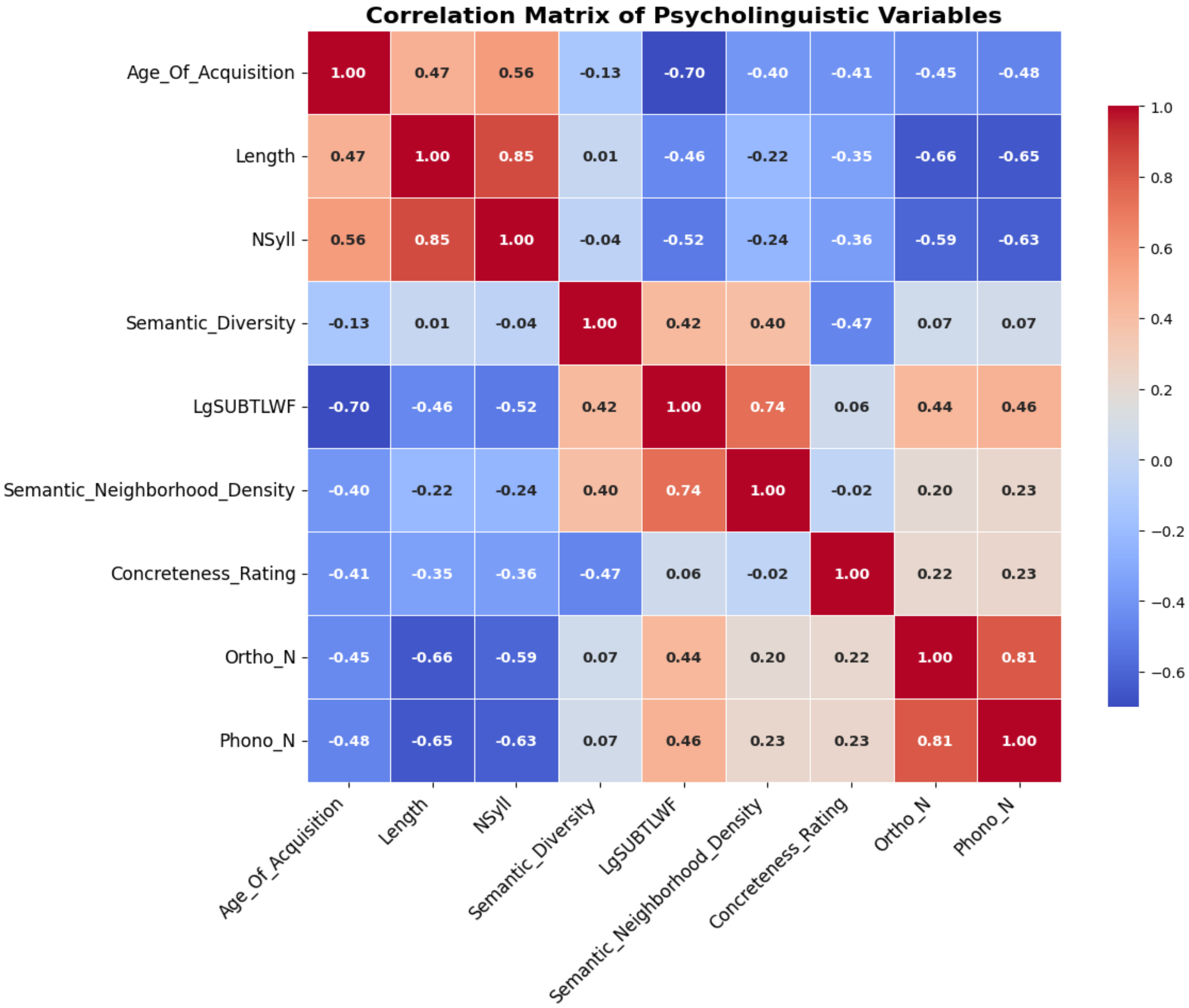


Pairwise Pearson correlation coefficients are displayed for all variables. Correlation coefficients are unitless and range from −1 to +1, where positive values indicate a positive linear association, negative values indicate a negative linear association, and values closer to zero indicate weaker or no linear relationship. The matrix is visualised using a diverging blue-red colour scheme, with dark blue representing strong negative correlations (−1), white representing correlations close to zero, and dark red representing strong positive correlations (+1). The colour bar on the right indicates the mapping of correlation coefficients to colours. Numerical correlation coefficients are overlaid within each cell (rounded to two decimal places), and the main diagonal contains values of 1, representing each variable's correlation with itself.

*Nyll = Number of syllables; LgSUBTLWF = log10 transform of the SUBTLEX frequency per million words; Ortho_N = number of orthographic neighbours; Phono_N = number of phonological neighbours (excluding homophones). Definitions and details of all psycholinguistic measures included in the PCA are available at the English Lexicon Project: [https://elexicon.wustl.edu](https://elexicon.wustl.edu).*

Supplementary Materials 4: *SJT LMEMs and the effect of psycholinguistic properties on accuracy and response time.*

*The top row of graphs depict the effect of PCA factors on the predicted probability of response time (ms) and the bottom row depict their effect on the predicted probability of accuracy. The x axes are consistent across panels and indicate the relevant factor increasing in score with descriptions in brackets to aid in the interpretation of the direction of each x axis. The first column pertains to Factor 1 (Word-Form Simplicity), the second column to Factor 2 (Semantic Abstractness), and the third column to Factor 3 (Lexical Familiarity).*

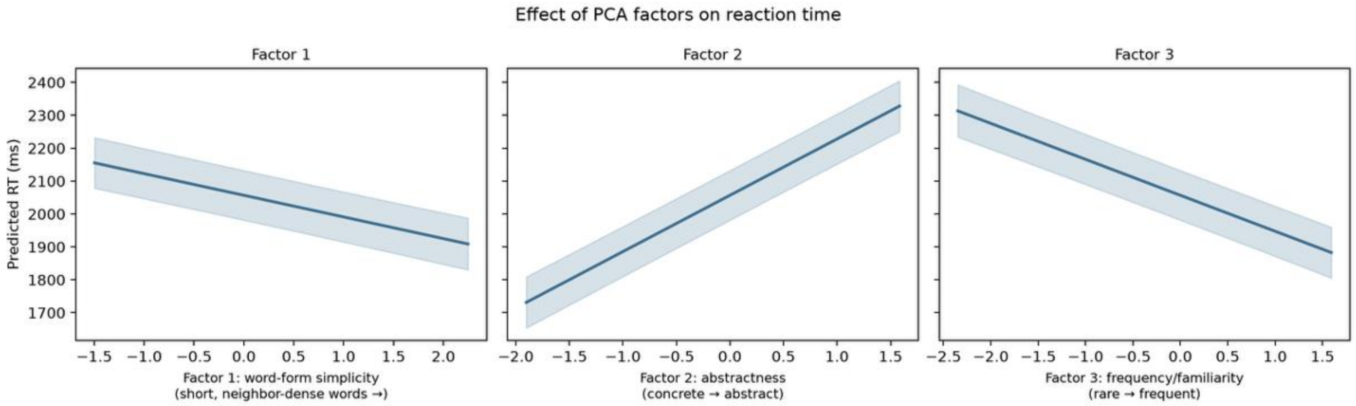


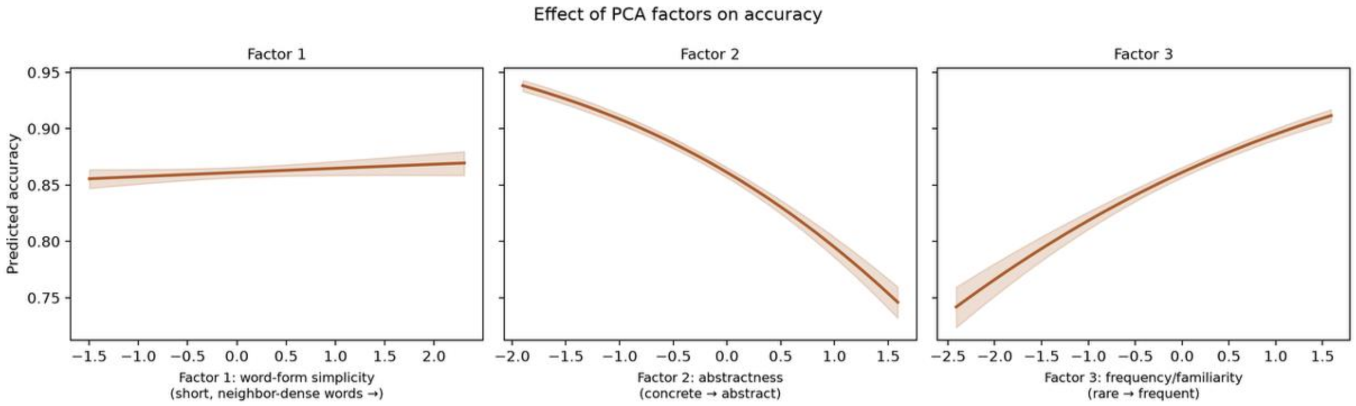

Supplementary Materials 5: *VGT LMEMs and the effect of five manipulated perceptual properties on accuracy and reaction time.*

*Graphs in panel (a) (top row, green) depict the predicted probability of accuracy (0.5 – 1.0) based on Grid Dimension, Number of Colours, Colour Ratio, Rotation, and Flip (L-R). Graphs in panel (b) (bottom row, blue) depict the predicted probability of response time (ms; 2000 - 2750) based on these same variables. Both y axes (in the accuracy and response time rows) are consistent across all panels within a row for comparison.*

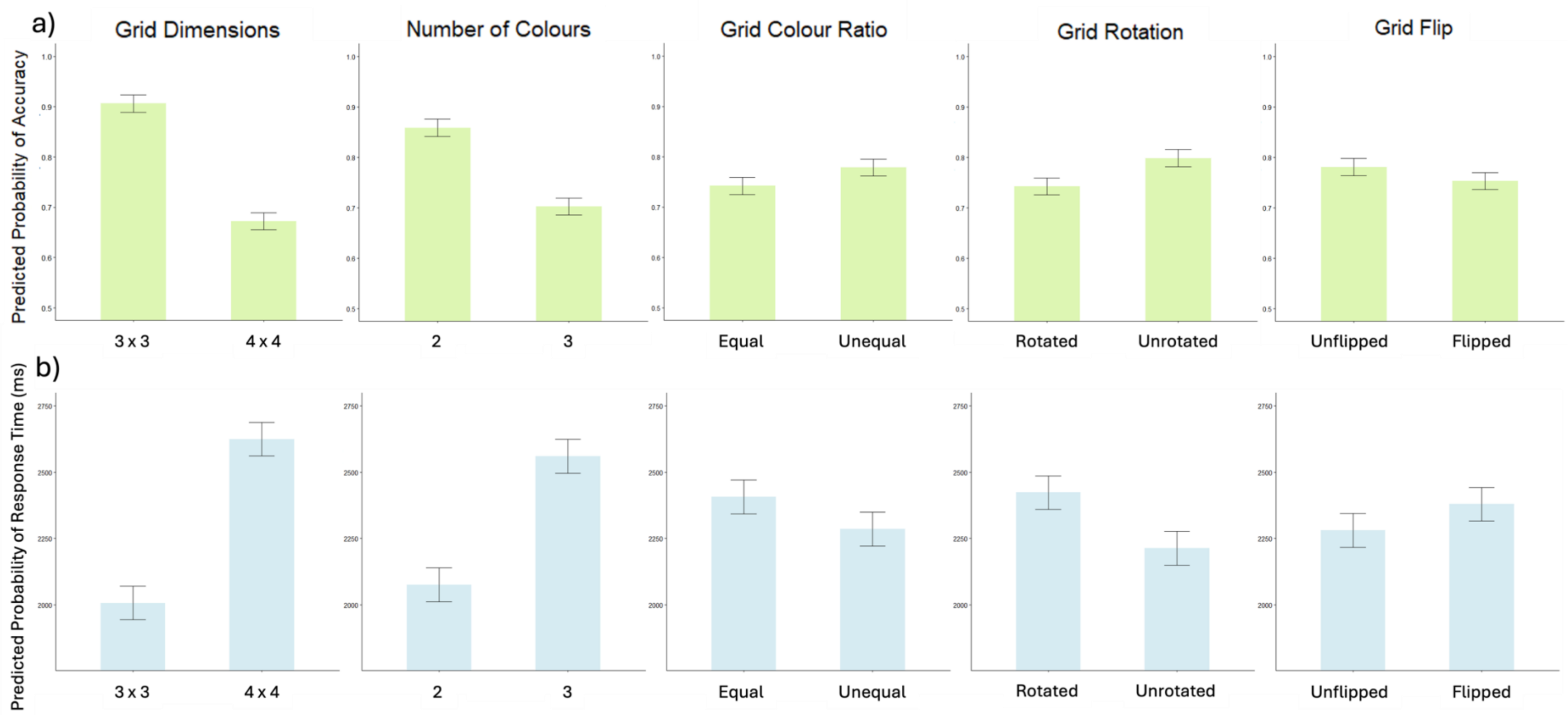
